# Inflammatory restraint and membrane lipid integrity protect hematopoietic stem cells under stress

**DOI:** 10.64898/2026.08.24.746871

**Authors:** Ayano Yahagi, Haruka Okabe-Kitajima, Makiko Mochizuki-Kashio, Kyoko Komai, Takayoshi Matsumura, Terumasa Umemoto, Mikiro Nawa, Fumio Nakamura, Takayuki Yoshimoto, Kohsuke Kanekura, Stephanie Z. Xie, Keiyo Takubo, Toshio Suda, Ayako Nakamura-Ishizu

## Abstract

Life-long production of blood requires the preservation of hematopoietic stem cell (HSCs) regenerative capacity during inflammation. The cytokine, Thrombopoietin (THPO), is essential for HSC maintenance yet its role during inflammatory stress remains incompletely understood. Long-term repopulating potential was rapidly depleted in THPO-deficient HSCs upon poly(I:C) administration through inflammatory pyroptosis. Transcriptomic and chromatin accessibility analyses revealed constitutive interferon (IFN) pathway activation in THPO-deficient HSCs, characterized by enhanced STAT1 signaling, increased accessibility of STAT and IRF motifs, and elevated expression of IFN-stimulated genes. Lipidomic profiling further identified selective shifts in sphingomyelin (SM) species and enrichment of features associated with increased bilayer rigidity. THPO-deficient HSCs displayed elevated membrane SM incorporation, impaired membrane fluidity and altered membrane ultrastructure. Genetic ablation of *Stat1* normalized membrane lipid abnormalities and reduced pyroptotic activation and restored HSC survival and regenerative function under inflammatory stress. Together, these findings identify a STAT1 and SM metabolism as critical THPO downstream to protect HSCs from inflammatory pyroptosis. Our results reveal membrane lipid homeostasis as a fundamental mechanism through which cytokine signaling safeguards HSC function during stress.

## Introduction

Hematopoietic stem cells (HSCs) sustain lifelong blood production through a tightly regulated balance between quiescence, self-renewal and differentiation ^1–4^. Maintenance of stem-cell integrity is particularly critical during inflammatory stress as excessive or chronic inflammatory signaling compromises stem-cell function, promotes exhaustion and contributes to aging and disease ^5–9^.

Apart from regulating megakaryopoiesis, THPO and its receptor MPL are required for bone marrow (BM) HSC self-renewal and maintenance ^10–13^. THPO-and MPL-deficient mice exhibit an age-dependent decline in the BM HSC numbers ^13,14^ and notably defective protection to DNA damage upon replication in HSCs ^15,16^. Yet despite reductions in BM HSC frequency at steady-state, we report that HSCs from THPO-deficient mice demonstrate intact repopulation potential when transplanted to wild-type mice ^17,18^, which questions whether THPO functions are limited to achieving and maintaining a substantial number of HSCs within the BM rather than affecting the stem cell potential of individual HSCs.

Microbial-induced inflammation involves the activation of IFN pathway that is essential for coordinating innate and adaptive immune responses ^19,20^. Through activation of JAK/STAT pathways, IFN signaling trains and primes immune responses of hematopoietic cells by influencing epigenetic and cell metabolic states ^20–22^. Transient IFN signaling can activate dormant HSCs, persistent activation of IFN-responsive pathways is associated with impaired stem-cell function as has been demonstrated through analyzing mouse systemic infection models using the viral mimetic polyinosinic:polycytidylic acid (poly(I:C)) ^5,6,23^. Excessive IFN signaling also drives acute transient skewed differentiation of HSCs to megakaryocytes ^24^. While THPO/MPL signaling involves JAK/STAT activation similar to IFN signaling, the mechanisms underlying the differing consequential signal-induced cell fates remain an enigma. Furthermore, whether and how THPO/MPL is required to protect HSCs from inflammatory stress and the molecular pathways underlying such protection remain poorly understood.

Here we show that THPO protects HSCs from inflammation by simultaneously restraining tonic STAT1 signaling and maintaining cell membrane lipid homeostasis. In the absence of THPO, HSCs acquire a constitutively IFN-primed state characterized by enhanced STAT1 activity and remodeling of membrane SM composition. These alterations reduce membrane fluidity and lower the threshold for inflammatory pyroptosis. Mechanistically, genetic inhibition of STAT1 partially rescued HSC survival and regenerative function under inflammatory stress. Our findings uncover an unexpected link between cytokine signaling, membrane lipid biology and stem-cell survival and identify membrane integrity as a critical determinant of HSC stress tolerance.

## Results

### THPO prevents HSCs from inflammatory pyroptosis

HSCs from *Thpo^-/-^*mice under no stress exhibit similar repopulation potential compared to HSCs from *Thpo^+/+^* mice ^17,25^ but less is known how THPO deficiency affects HSC stem cell potential during inflammatory stress. We thus administered poly(I:C), a synthetic double-stranded RNA analog that induces systemic inflammatory signaling to *Thpo^+/+^*and *Thpo^-/-^* mice (Fig. 1A). After 48hrs of poly(I:C) administration, *Thpo^-/-^* HSCs exhibited significant reduction in both frequency and absolute number compared to *Thpo^+/+^* HSCs, indicating heightened susceptibility to inflammatory stress-induced depletion (Fig. 1B,C). Serial competitive BM transplantation revealed a striking decrease of PB chimerism in both myeloid and lymphoid lineages for HSCs from *Thpo^-/-^* mice treated with poly(I:C) compared to HSCs from *Thpo^+/+^* mice treated with or without poly(I:C) (Fig.1D). BM hematopoietic stem and progenitor (HSPC) subsets also showed significantly low chimerism in recipients transplanted with HSCs from *Thpo^-/-^* mice treated with poly(I:C) compared to HSCs from *Thpo^+/+^* mice treated with or without poly(I:C) in both primary and secondary transplantation (Fig.1E).

**Figure 1:**
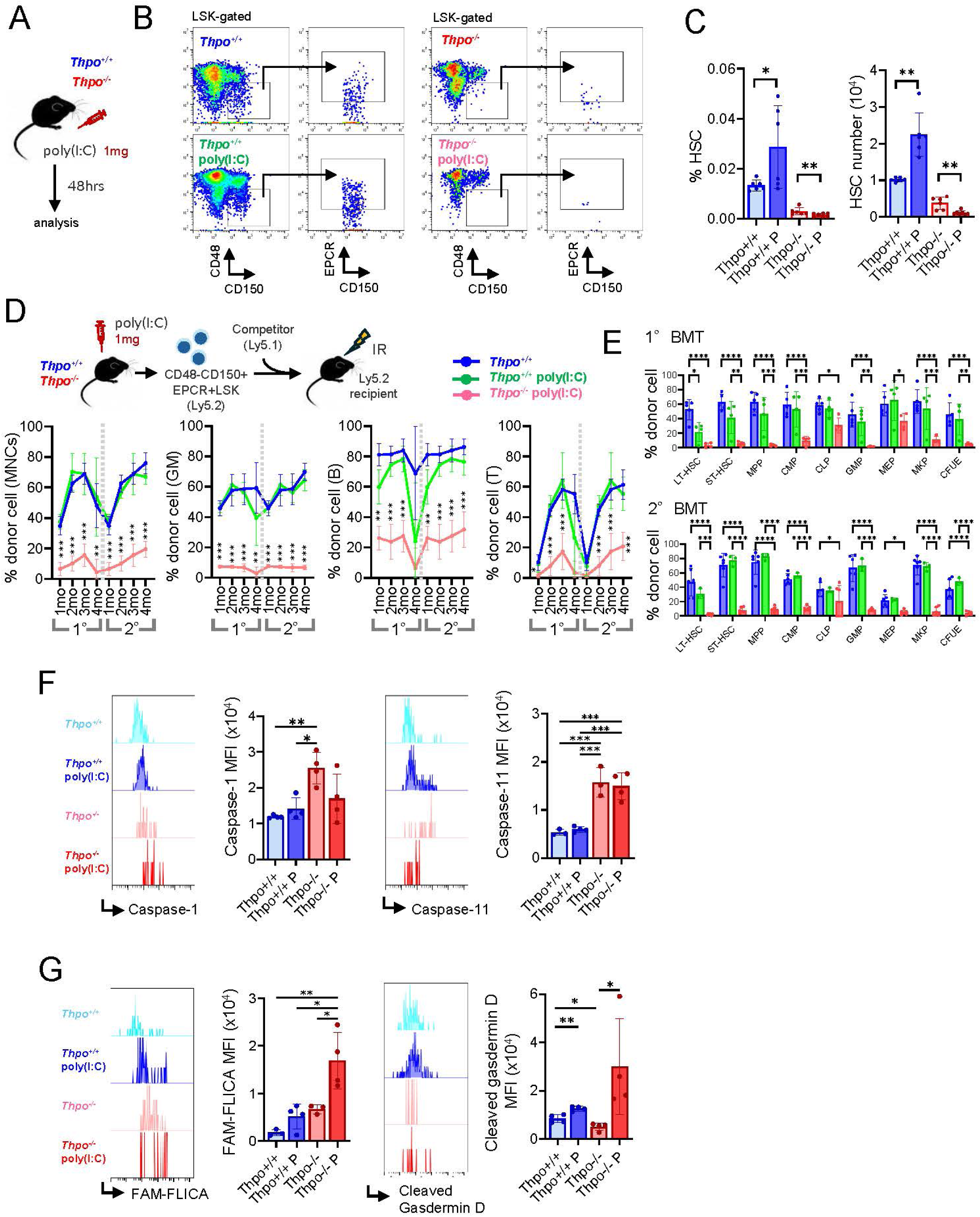
THPO-deficient HSCs rapidly lose stem cell potential through pyroptosis during inflammatory stress (A) Schematic representation of poly(I:C) administration to *Thpo^+/+^* and *Thpo^-/-^* mice. (B) Representative flow cytometric plots of HSC (CD48-, CD150+, EPCR+ LSK cells) from *Thpo^+/+^* and *Thpo^-/-^*mice with or without poly(I:C) administration. (C) HSC frequency and number from *Thpo+/+* and *Thpo^-/-^* mice with or without poly(I:C) (P) administration. (mean±SD) (n=6) * P<0.05, ** P<0.01 by Tukey test. (D) Primary competitive BM transplantation of 500 HSCs isolated from non-treated *Thpo^+/+^*mice and *Thpo^+/+^* and *Thpo^-/-^* mice treated with poly(I:C). Monthly PB chimerism of total MNCs, GM cells (Mac1+, Gr1+), B cells (B220+) and T cells (CD4+, CD8+) for primary and secondary BM transplantation. (mean±SD) (n>4) ** P<0.01, *** P<0.001 by Tukey test. (E) BM chimerism of LT-HSCs (CD48-, CD150+, EPCR+ LSK cells), ST-HSCs (CD34+, Flt3-LSK cells), MPPs (CD34+, Flt3+ LSK cells), CMPs (CD16/32-CD34+ Lin-cKit+Sca1-(LKS-) cells), CLPs (CD16/32+, Flt3+ LKS-cells), GMPs (CD16/32+ LKS-cells), MEPs (CD16/32-, CD34-LKS-cells), MkPs (CD41+, CD150+ LKS-cells) and CFUEs (CD16/32-, CD150-, CD105+, Ter119-LKS-cells) (mean±SD) (n>4) * P<0.05, ** P<0.01, *** P<0.001 and **** P<0.0001 by Tukey test. (F) Representative flow cytometric plots and quantification (mean fluorescence intensity (MFI)) of caspase-1 and caspase-11 on HSCs from *Thpo^+/+^* and *Thpo^-/-^*mice with or without poly(I:C) (P). (mean±SD) (n>3) * P<0.05, ** P<0.01 and *** P<0.001 by Tukey test. (G) Representative flow cytometric plots and MFI of active caspase-1 (FAM-FLICA) and cleaved-gasdermin D on HSCs from *Thpo^+/+^* and *Thpo^-/-^* mice with or without poly(I:C) (P). (mean±SD) (n>3) * P<0.05 and ** P<0.01 by Tukey test.

Given the rapid loss of HSC number and stem cell potential, activation of inflammatory cell death pathways were analyzed. THPO-deficient HSCs exhibited significantly high levels of caspase-1 and caspase-11 prior and following poly(I:C) treatment indicating inflammasome activation and pyroptotic execution (Fig. 1F). Furthermore, THPO-deficient HSCs exhibited a significant upregulation of activated caspase-1 (FAM-FLICA staining) and cleaved-gasdermin D with poly(I:C) treatment indicating execution of pyroptosis (Fig. 1G). In contrast, THPO-deficient HSCs exhibited downregulation of Annexin V staining with poly(I:C) treatment (Extended data 1A). Increase in caspase-1 and -11 was also evident in HSCs deficient of *Mpl*, the receptor of THPO (Extended data 1B,C). Moreover, although BM HSC number and frequency was intact (Extended data 1D,E), lipopolysaccharide (LPS) administration stimulated pyroptosis in *Thpo^-/-^*HSCs (Extended data 1F). These findings indicate that THPO protects HSCs from inflammatory pyroptosis and preserves stem cell function under inflammatory stress.

### THPO deficiency induces an interferon-responsive STAT1 active state in HSCs

To understand the molecular adaptations underlying HSC response to inflammatory stress in the absence of THPO, we re-examined our single cell RNA sequence (scRNAseq) data of HSPCs from *Thpo^+/+^* and *Thpo^-/-^* mice ^25^. Unsupervised clustering resolved two major HSC clusters (HSC-1 and HSC-2) (Extended data 2A), revealing a substantial reduction of HSC-1 cluster in *Thpo^-/-^*mice (Fig. 2A). Gene set enrichment analysis (GSEA) identified strong activation of IFN-associated programs, including IFNγ response, antiviral defense pathways and STAT1 target gene signatures, in HSC-2 cluster cells that were observed predominantly in *Thpo^-/-^*(Fig. 2B).

**Figure 2:**
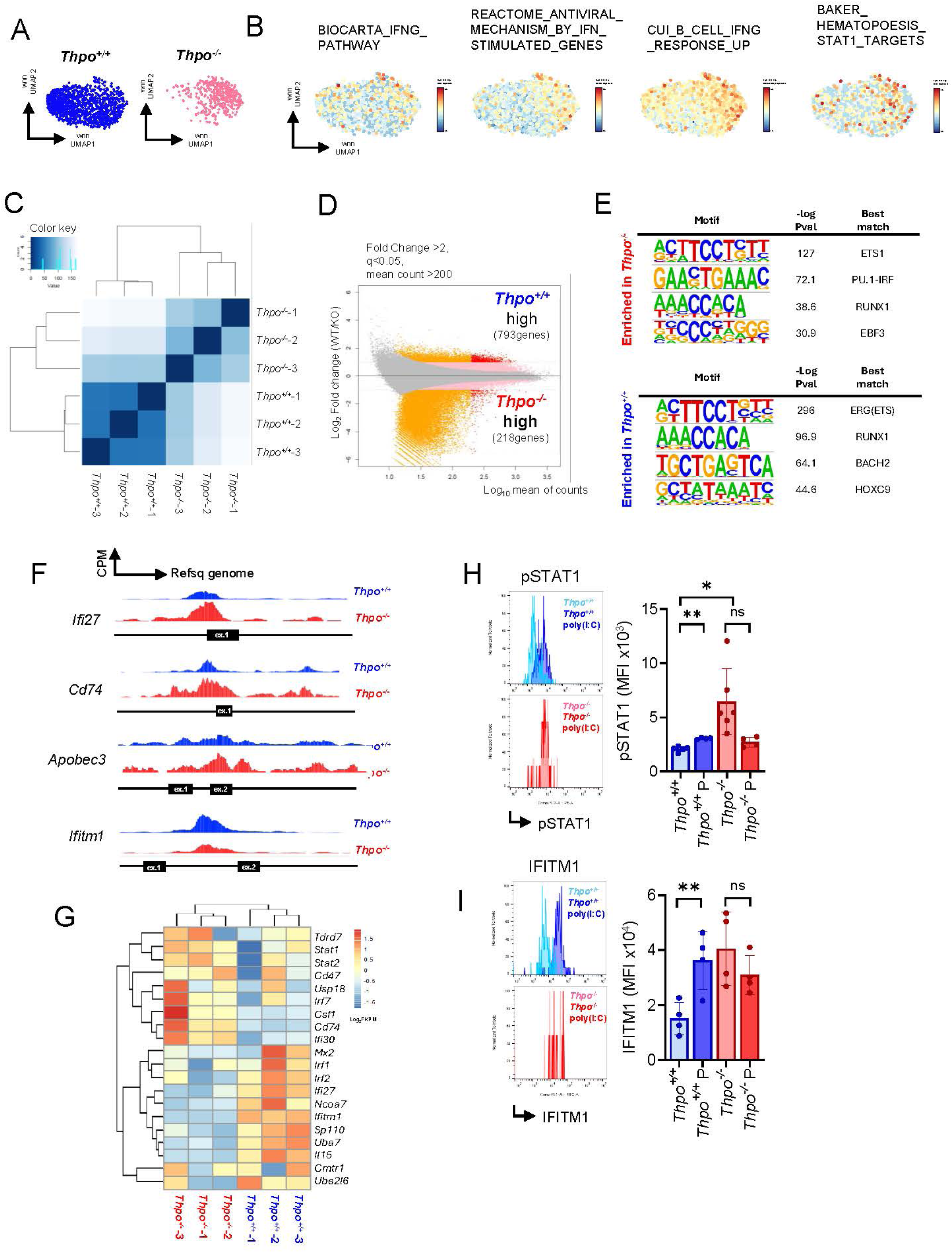
THPO deficiency induces an interferon-responsive STAT1 active state in HSCs (A) UMAP of single-cell RNA sequence data of HSCs from *Thpo^+/+^* and *Thpo^-/-^*mice. (B) Expression of various IFN-related gene sets in HSC cluster. Note that *Thpo^-/-^*HSC dominant cluster exhibits a higher enrichment of these gene sets. (C) Hierarchical clustered heatmap from ATAC-seq signals for *Thpo^+/+^* and *Thpo^-/-^* HSCs. (D) MA plot of differential chromatin accessibility for values with fold Change >2, q-val<0.05 andmean count >200 for *Thpo^+/+^* and *Thpo^-/-^*HSCs. (E) Homer motif analysis for motifs enriched in *Thpo^+/+^* or *Thpo^-/-^*HSCs. (F) Counts per million (CPM) plots on reference genome for *Ifi27, Cd74, Apobec3* and *Ifitm1* comparing open chromatin regions between *Thpo^+/+^* and *Thpo^-/-^*HSCs. (G) Expression of IFN-associated genes in bulk RNA-seq data of *Thpo^+/+^*and *Thpo^-/-^*HSCs. (H)(I) Representative flow cytometric plots and quantification of pSTAT1 and IFITM1 expression on HSCs from *Thpo^+/+^*and *Thpo^-/-^* mice treated with poly(I:C). (mean±SD) (n>4) * P<0.05, ** P<0.01, ns: not significant by Tukey test.

To determine whether this transcriptional adaptation was associated with epigenetic remodeling, we performed ATAC-seq on purified HSCs from *Thpo^+/+^* and *Thpo^-/-^*mice. Compared to *Thpo^+/+^* HSCs, *Thpo^-/-^* HSCs exhibited changes in chromosome accessibility with over 30% locating in promotor (TSS±3kb) regions (Fig. 2C, Extended data 2B,C). Further analysis narrowing data to more significantly open peaks (Fold Change >2, q-val<0.05 and mean count >200) revealed enriched accessibility in 793 genes for *Thpo^+/+^* HSCs and 218 genes for *Thpo^-/-^* HSCs (Extended data 2D, Fig. 2D). Motif enrichment analysis revealed high accessibility at interferon regulatory factor (PU.1-IRF) motif in *Thpo^-/-^* HSCs compared to *Thpo^+/+^*HSCs (Fig. 2E). Gene ontology analysis also exhibited an enrichment in gene sets associated with immune and inflammatory response in *Thpo^-/-^* HSCs (Extended data 2E). Among the 218 genes enriched in *Thpo^-/-^* HSCs, we found canonical IFN-stimulated genes (ISGs) (*Ifi27, Cd74, Apobec2*) and genes related to IFN signaling (*Ifi30, Znfx1, H2-DMb2, H2-Eb2*) with notably increased accessibly near TSS sites (Fig. 2F, Supplementary file 1). On the other hand, *Ifitm1* was the only ISG among the 793 genes that were enriched in *Thpo^+/+^* HSCs (Fig. 2F, Supplementary file 1). Based on these epigenetic changes, we re-examined our bulk RNA-seq data conducted on *Thpo^+/+^*and Thpo*^-/-^* HSCs ^17^, which indicated an increase in the expression of ISGs identified in the ATAC-seq (*Ifi27, Cd74, Ifi30*) as well as other ISGs including *Stat1* and *Stat2* in *Thpo^-/-^* HSCs (Fig. 2G). However, expression of canonical ISGs such as *Mx2, Ifitm1, Sp110* and *Uba7* were high in *Thpo^+/+^* HSCs compared to *Thpo^-/-^*HSCs (Fig. 2G).

Taken the variations in epigenetic accessibility and genetic expressions of IFN-associated genes, we analyzed protein expressions of phosphorylated-STAT1 (pSTAT1) and IFITM1, a critical ISG product expressed on the cellular membrane ^26^ (Fig. 2H). Compared to *Thpo^+/+^* HSCs, *Thpo^-/-^* HSCs exhibited significantly high expression of pSTAT1 and both cytoplasmic and membrane-bound IFTIM1 (Fig. 2H, Extended data 3). While pSTAT1 expression was upregulated in *Thpo^+/+^* HSCs and suppressed in *Thpo^-/-^* HSCs, IFITM1 expression was upregulated in *Thpo^+/+^* HSCs and remained high in *Thpo^-/-^*HSCs upon poly(I:C) administration (Fig. 2I). Convergent scRNA-seq and ATAC-seq data with increased pSTAT1 and IFITM1 protein expression support a constitutive STAT1-high state in THPO-deficient HSCs. Together, these findings reveal that THPO deficiency establishes a state of persistent IFN-signaling that primes HSCs to undergo pyroptotic cell death upon inflammation.

### THPO maintains membrane sphingolipid homeostasis and preserves HSC membrane fluidity

Given the constitutive inflammatory activation observed in *Thpo^-/-^*HSCs, we next asked how this altered inflammatory state made THPO deficient HSCs more vulnerable to pyroptosis. Multiple ISGs including IFITM1 are known to influence cell membrane integrity to restrict viral entry during inflammation ^27,28^. Moreover, lipid metabolic remodeling of the plasma membrane has been implicated in inflammatory stress response ^29–31^. Unbiased lipidomic profiling of LSK cells from *Thpo^+/+^* and *Thpo^-/-^* mice detected 390 lipid species from 14 lipid subclasses. Principal component analysis (PCA) of lipidomics data exhibited segregation between the lipid profile of *Thpo^-/-^*and *Thpo^+/+^* HSPCs indicating broad lipid compositional remodeling in the absence of THPO (Fig. 3A). Functional annotations of altered lipid species using LION enrichment analysis ^32^ revealed enrichment of membrane biophysical features associated with reduced lateral diffusion, increased bilayer rigidity and altered membrane curvature indicative of increased cell membrane rigidity (Fig. 3B, C). Expressions of membrane-associated lipids (sphingomyelin (SM), phosphatidylserine (PS), phosphatidylinositol (PI), phosphatidylethanolamine (PE), phosphatidylcholine (PC) and ceramide (Cer)) were prominently altered with THPO deficiency (Extended data 4A). While the direct quantification of total cellular SM, which is known to increase cell membrane rigidity ^33,34^, was comparable between *Thpo^+/+^* and *Thpo^-/-^* HSCs, across multiple SM species, *Thpo^-/-^*HSCs exhibited increase in SM that are more saturated (SM(d34:1), SM(d34:0) and with long fatty acid chains (SM(d42:2)) which may increase lipid membrane order ^35^ (Extended data 4B,C). Transcriptional analysis of sphingolipid metabolic enzymes revealed upregulation of expression of genes involved in SM synthesis (*Sgms2*), down regulation of expression of genes involved in SM degradation (*Smdpl3a, Smdpl3b, Smpd2, Smpd3* and *Smpd5)* and down regulation of expression of genes involved in ceramide degradation (*Asah2, Adipor2* and *Acer2)* in *Thpo^-/-^* HSCs (Fig. 3D). Moreover, ATAC-seq data showed that *Thpo^-/-^* HSCs exhibited an enrichment in accessibility of membrane lipids and sphingolipid associated genes (Extended data 4D).

**Figure 3:**
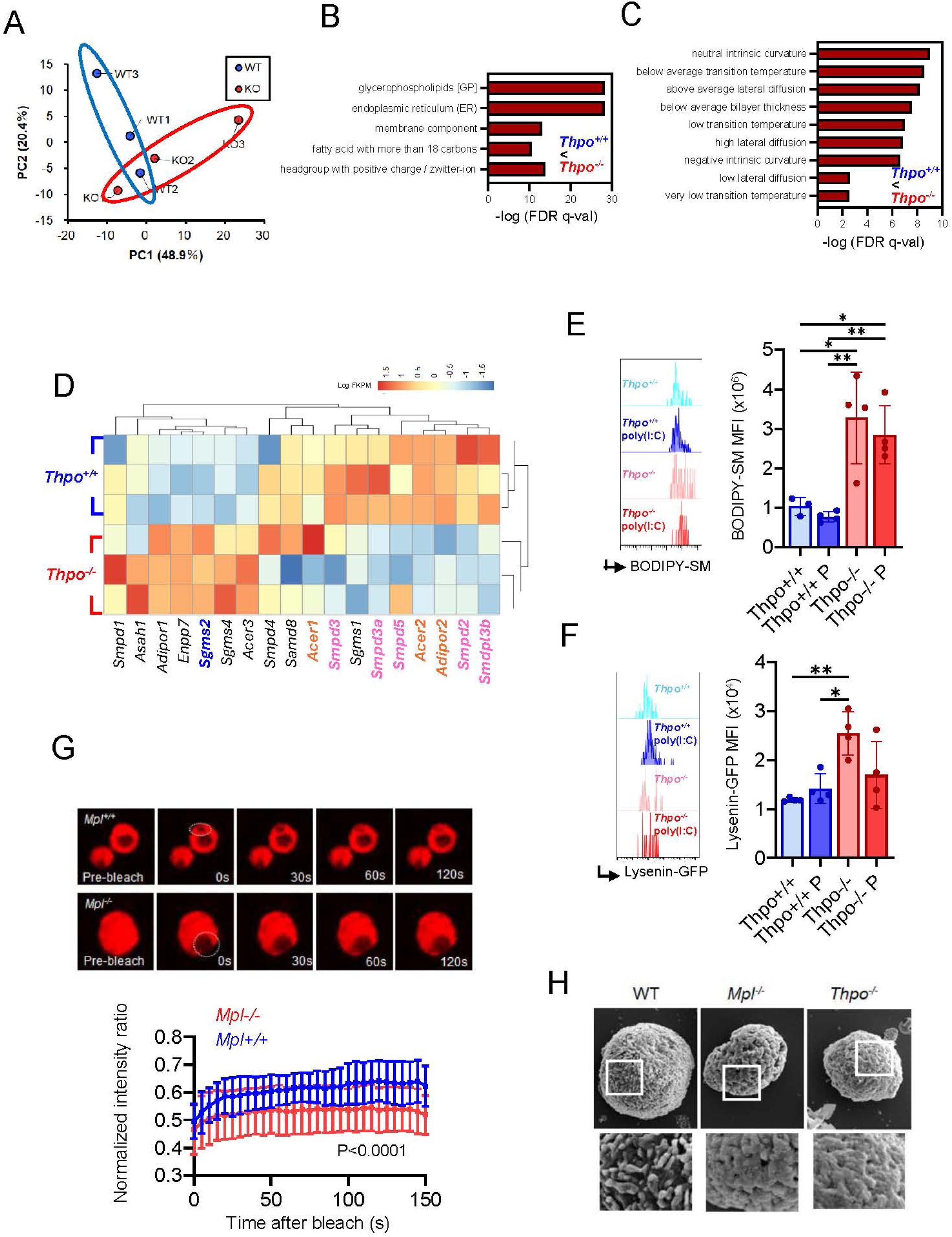
THPO protects HSCs from inflammatory pyroptosis through regulating cell membrane integrity (A) Principal component analysis (PCA) plot for lipidome analysis of *Thpo^+/+^*(WT) and *Thpo^-/-^* (KO) HSCs. (n=3) (B) Lipid ontology (LION) enrichment analysis showing -log (FDR q-val) for lipid sets that are enriched higher in *Thpo^-/-^* HSCs compared *Thpo^+/+^*HSCs. (C) LION analysis showing -log (FDR q-val) for membrane-associated lipid sets that are enriched higher in *Thpo^-/-^* HSCs compared *Thpo^+/+^* HSCs. (D) Representative flow cytometric plot and quantification of BODIPY-SM staining on cell membrane in HSCs from *Thpo^+/+^* and *Thpo^-/-^*mice treated with poly(I:C). (mean±SD) (n>3) * P<0.05 by Tukey test. (E) Representative flow cytometric plot and quantification of lysenin-GFP staining on cell membrane in HSCs from *Thpo^+/+^* and *Thpo^-/-^* mice treated with poly(I:C). (mean±SD) (n>3) * P<0.05, ** P<0.01 by Tukey test. (F) Heatmap showing gene expressions from bulk-RNAsq of various genes coding factors associated with SM in HSCs from *Thpo^+/+^* and *Thpo^-/-^*mice. Note that genes involved in SM synthesis are indicated in blue, genes involved in SM degradation are indicated in pink and genes involved in ceramide degradation are indicated in orange. (G) Representative images showing fluorescence recovery after photobleaching (FRAP) analysis on *Mpl^+/+^* and *Mpl^-/-^* and WT HSCs cultured without THPO. Normalized intensity ratio at indicated time after bleaching using FRAP. P<0.0001 by non-linear regression (curve fit) one phase association. (H) Scanning electron microscopic images of WT, *Mpl^-/-^*and *Thpo^-/-^* HSCs.

We next quantified the ability of HSC cell membrane to incorporate SM using a BODIPY-labelled SM probe. Under our staining conditions (optimized staining at 4℃ for 20min, Extended data 4E), BODIPY–SM reports probe incorporation into the cellular membrane and serves as a surrogate for cell membrane-accessible SM. HSCs from *Thpo^-/-^* and *Mpl^-/-^* mice exhibited increased membrane-associated SM at baseline, which was further enhanced following poly(I:C) challenge (Fig. 3E, Extended data 4E,F). Furthermore, direct quantification of cell membrane SM by staining with GFP-conjugated lysenin ^36^, a protein that binds directly to SM, showed significant upregulation of membrane SM in HSCs from *Thpo^-/-^* and *Mpl^-/-^* mice (Fig 3F, Extended data 4G). These findings indicate that THPO deficiency may induce persistent accumulation of membrane SM under both homeostatic and inflammatory conditions which may influence membrane rupture susceptibility in HSCs during inflammatory stress.

To directly assess the functional consequences of altered membrane lipid composition, we measured membrane dynamics using fluorescence recovery after photobleaching (FRAP). *Mpl*-deficient HSCs exhibited significantly impaired fluorescence recovery kinetics, indicating reduced membrane fluidity and diminished lateral mobility within the plasma membrane (Fig. 3G, Extended data video A,B). These data establish that THPO signalling actively preserves membrane fluidity. Scanning electron microscopy (SEM) images of the cell surface exhibited revealed less ridges and grooves present on *Mpl^-/-^*and *Thpo^-/-^* HSCs compared to wild-type (WT) cells (Fig. 3H). Together, these findings identify THPO as a regulator of HSC membrane sphingolipid homeostasis and suggest that THPO deficiency promotes cell membrane SM enrichment and reduced fluidity in HSCs, which permits inflammatory pyroptotic membrane rupture.

### STAT1 deletion restores HSC resistance to inflammatory pyroptosis

To determine whether elevated STAT1 signaling directly contributes to the inflammatory vulnerability of THPO-deficient HSCs, we generated compound mutant mice lacking both *Thpo* and *Stat1* (*Thpo^-/-^;Stat1^-/-^*) and assessed pyroptotic activation following poly(I:C) challenge (Fig. 4A). While STAT1 deficiency alone did not affect pyroptotic processes at steady state (Extended data 5A), *Stat1^-/-^* HSCs exhibited decreased staining towards BODIPY-SM but not lysenin-GFP (Extended data 5B). Combined deletion of *Thpo^-/-^*and *Stat1^-/-^* exhibited that *Stat1-*deletion slightly but not significantly upregulated BM HSC number in *Thpo^-/-^*mice upon poly(I:C) administration (Fig. 4B). Yet, *Thpo^-/-^;Stat1^-/-^*HSCs exhibited significantly reduced caspase activity and cleaved gasdermin-D to a level comparable to that of wild-type and *Stat1-/-* controls (Fig. 4C,D). Furthermore, *Stat1-*deletion significantly down-regulated staining intensity of BODIPY-SM and lysenin-GFP in THPO- deficient HSCs indicating that STAT1 functions upstream of membrane lipid remodeling (Fig. 4E,F). Furthermore, HSCs from *Thpo^-/-^;Stat1^-/-^*mice treated with poly(I:C) exhibited an increase in fluorescence recovery after bleach, indicating a recovery of cell membrane fluidity (Fig. 4G,H) These findings demonstrate that STAT1-deletion partially rescues pyroptotic susceptibility associated with THPO- deficiency. Together, THPO/MPL signaling suppresses aberrant STAT1 activation, maintains membrane lipid homeostasis and membrane integrity, thereby protecting HSCs from inflammatory pyroptotic collapse (Fig 4I).

**Figure 4:**
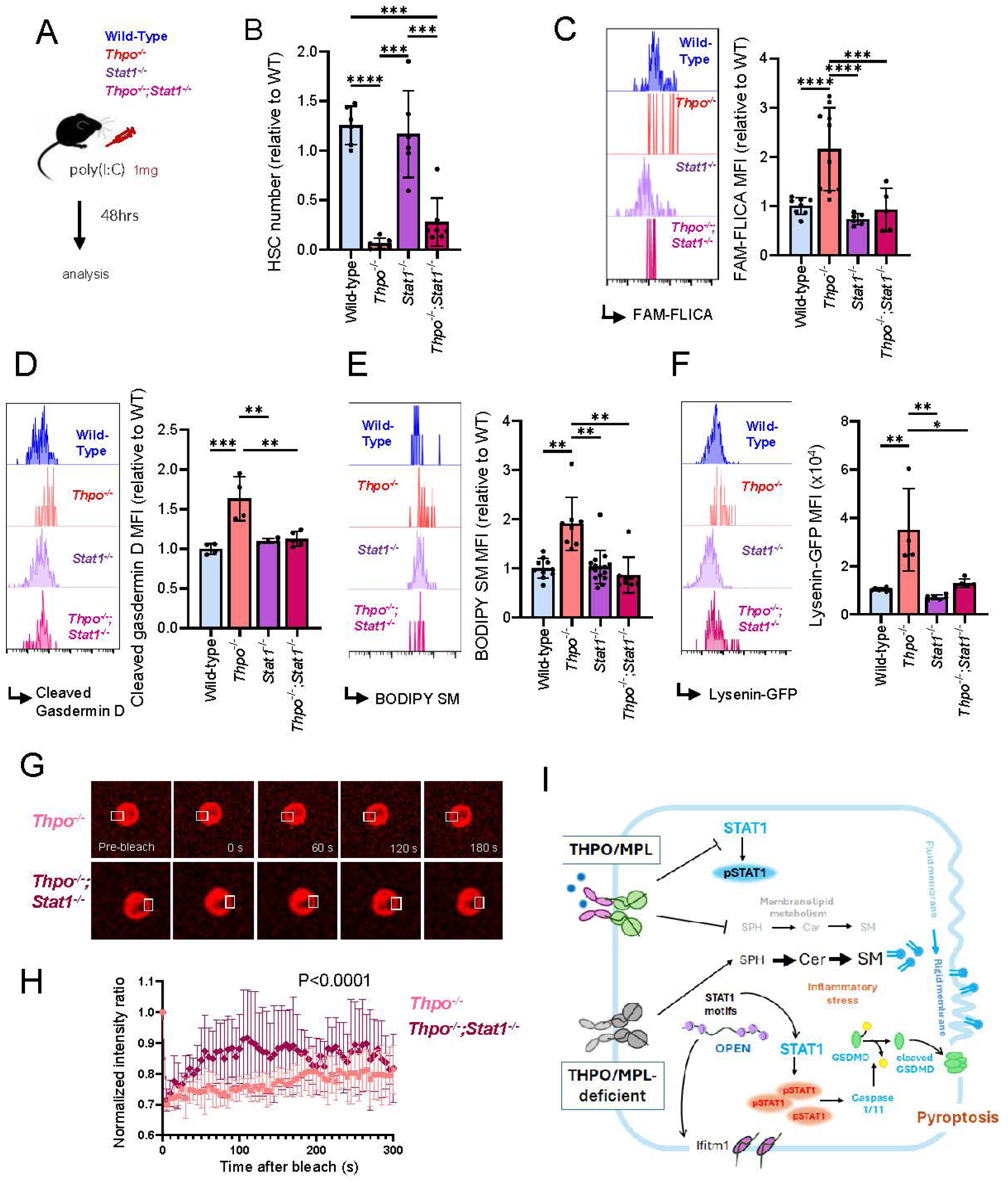
STAT1 depletion ameliorates inflammatory pyroptosis in HSCs (A) Schematic representation of poly(I:C) administration to wild-type (WT), *Thpo^-/-^*, *Stat1^-/-^* and *Thpo^-/-^;Stat1^-/-^* mice. (B) HSC number of WT, *Thpo^-/-^*, *Stat1^-/-^* and *Thpo^-/-^;Stat1^-/-^*mice treated with poly(I:C). Values are shown relative to WT. (mean±SD) (n>5) *** P<0.001 and **** P<0.0001 by Tukey test. (C-F) Representative flow cytometric plots and MFI of (C) active caspase-1 (FAM-FLICA), (D) cleaved-gasdermin D, (E) BODIPY-SM and (F) Lysenin-GFP staining in HSCs from WT, *Thpo^-/-^*, *Stat1^-/-^*and *Thpo^-/-^;Stat1^-/-^* mice treated with poly(I:C). MFI values are shown relative to WT. (mean±SD) (n>4) * P<0.05, ** P<0.01, *** P<0.001 and **** P<0.0001 by Tukey test. (G) Representative images showing fluorescence recovery after photobleaching (FRAP) analysis on HSCs from *Thpo^-/-^* and *Thpo^-/-^; Stat1^-/-^* mice treated with poly(I:C). (H) Normalized intensity ratio at indicated times after bleaching using FRAP. P<0.0001 by non-linear regression (curve fit) one phase association. (I) Graphic image illustrating the effect of THPO/MPL signaling during inflammatory stress.

## Discussion

Our study identifies a previously unrecognized mechanism through which THPO/MPL signaling safeguards HSC integrity during inflammatory stress. We demonstrate that THPO suppresses constitutive IFN–STAT1 signaling and maintains membrane sphingolipid homeostasis, thereby preserving membrane fluidity and protecting HSCs from pyroptotic cell death. Loss of THPO establishes a maladaptive state characterized by persistent inflammatory priming, SM accumulation and membrane rigidification, ultimately lowering the threshold for cell death during inflammatory stress.

THPO/MPL signaling reportedly is critical for the maintenance of HSCs during irradiation-induced replicative stress through modulation of DNA damage response and genomic integrity ^16,37^. Particularly, administration of THPO mimetic drugs rapidly upregulates STAT1 signaling which represses retroelements and mimics an anti-viral response to protect HSCs from genotoxic stress ^37^. On the contrary, our study identifies that germline depletion of THPO and MPL results in epigenetic and transcriptomic dysregulation of IFN-associated pathways with a prominent and consistent upregulation of STAT1 signaling in HSCs. We did not observe upregulation of pSTAT1 expression in adult conditional deletion of Thpo (*Thpo^fl/fl^;Cre^ERT^* mice ^25^) after 7 days of Cre-recombinase activation, suggesting that long-term THPO/MPL-deficiency that spans through developmental stages may rewire inflammatory downstream pathways in HSCs. The use of germline *Thpo^-/-^* and *Mpl^-/-^*models also cannot exclude the influence of changes in the HSC niche which may indirectly affect HSC cell fate. Megakaryocytes, which significantly decrease in number in *Thpo^-/-^*and *Mpl^-/-^* BMs, regulate HSCs ^38–40^. While the 48- hour timeframe between poly(I:C) administration and pyroptotic activation suggests acute molecular adaptation rather than chronic niche remodeling, the contributions of HSC-intrinsic versus niche-mediated mechanisms in establishing the IFN-primed, SM-rich state remain to be resolved through conditional deletion or reciprocal chimera approaches.

In our study, HSC vulnerability to inflammatory stress was partially rescued by STAT1 deletion. Loss of STAT1 minimally affects steady state, quiescent BM HSC numbers ^41^, yet *Stat1*-deficient mice are vulnerable to 5-fluorouracil-induced myeloablation due to decline in a specific HSC compartment that expressed high levels of MHCII ^42^. In the study, *Stat1*-deletion did not affect MHCII^low^ HSCs that were characterized as more proliferative and megakaryocyte lineage-biased, which are characteristics of HSCs that tend to rely more on THPO/MPL signaling ^43,44^. IFN-γ has also been reported to interfere with THPO binding to MPL^45^ whereas THPO receptor agonists such as eltrombopag may circumvent this interference and support human HSC survival and self-renewal ^46^. Collectively, these observations highlight the complex interplay between cytokine signaling and downstream effector activation and suggest that HSC maintenance may depend on a tightly regulated balance between inflammatory and THPO/MPL signaling. Maintaining this balance may be critical for protecting HSCs from inflammation-induced exhaustion and could provide a rationale for further exploring THPO receptor agonists in hematopoietic disorders and in the context of myeloablative conditioning for hematopoietic stem cell transplantation..

Emerging evidence indicates that lipid metabolism to inflammatory response of cells, especially the formation of inflammasomes that lead to pyroptotic cell death ^47^. A variety of stimuli may prime or activate NLRP3 (NACHT, LRR and PYD domains-containing protein 3) inflammasomes, including sphingosine and ceramide. While direct evidence lacks that high concentrations of SM associates with inflammasome activation, SM is associated with less fluid cell membranes susceptible to rupture and can mediate toxin binding to facilitate pore formation ^48,49^. While our study suggests that cytokines broadly regulate lipid metabolism, detailed analysis of whether and which SM metabolites can directly alter inflammasome activation should be further studied.

Collectively, we show that THPO/MPL signaling preserves HSC function under inflammatory stress through restraining inflammatory downstream signaling and maintaining cell membrane homeostasis. Our findings suggest that regulation of membrane lipid composition may represent a general mechanism by which tissue stem cells withstand inflammatory injury and maintain regenerative capacity throughout life.

## Author Contributions

A.Y. performed and analyzed some experiments and discussed the data. A.N-I designed the project, performed experiments, analyzed the data and wrote the manuscript. H.O-K performed and assisted in some of the experiments. T.M. performed and analyzed RNA-seq and ATAC-seq. M.N. and K.Kanekura. performed and assisted FRAP experiments and data analysis. T.Y. provided and assisted in analysis of *Stat1^-/-^* mice. M.M-K., K.Komai, T.U., F.N., S.X., K.T., and T.S. discussed and analyzed the data. All authors read and approved of the final manuscript.

## Acknowledgements

This study was supported by the KAKEN Grant-in-Aid for Scientific Research (B) (23K24366 to A.N-I), the KAKEN Grant-in-Aid for Scientific Research (C) (24K11298 to T.M.), JST FOREST (JPMJFR200G to A.N-I), AMED (24gm6710022h0001 to A.N-I) and grants from Astellas Foundation for Research on Metabolic Disorders (to A.N-I.), Daiichi Sankyo Foundation of Life Science (to T.M.), and the Uehara Memorial Foundation (to T.M.). SEM analysis was supported by Institute for Comprehensive Medical Sciences (ICMS), Tokyo Women’s Medical University.

## Materials and Method

### Animal models

All mice were on a C57BL/6N background and were 8-12 weeks of age. C57BL/6-Ly5.1 mice were used for competitive repopulation assays. *Thpo^-/-^*and *Mpl^-/-^* mice were previously described ^18,50^. *Stat1^-/-^* mice were obtained from Taconic and crossed with *Thpo^-/-^*mice to obtain *Thpo^-/-^;Stat1^-/-^* mice ^51^. Poly(I:C) and LPS administration were conducted as previously described ^24,52^. All the animal experiments were performed in accordance with the recommendations in the Guide for the Care and Use of Laboratory Animals of Animal Research Committee in Tokyo Women’s Medical University.

### Flow cytometric analysis

Flow cytometric analysis was performed as described previously (Nakamura-Ishizu et al., 2021). Briefly, suspensions of bone marrow (BM) cells from the femurs, tibiae, spine and iliac crest of C57BL/6NTac mice were isolated and depleted of red blood cells by an ammonium chloride solution. The following antibodies were used for flow cytometric analysis. c-Kit (2B8)(Biolegend), Sca-1 (D7)(Biolegend), CD4 (RM4-5)(Biolegend), CD8 (53-6.7) (Biolegend), B220 (RA3-6B2) (Biolegend), TER-119 (TER-119)(Biolegend), Gr-1 (RB6-8C5) (Biolegend), Mac-1 (M/70) (Biolegend), Flt-3 (A2F10)(Biolegend), CD34 (RAM34)(eBioscience), CD41 (MWReg30)(Biolegend), CD45.2 (104)(Biolegend), CD45.1 (A20) (Biolegend), CD16/32 (2.4G2)(BD Bioscience), CD48 (HM48-1)(Biolegend), CD150 (TC15-12F12.2)(Biolegend), IL-7Rα (A7R34)(eBioscience), Endoglin (MJ7/18)(Biolegend), AnnexinV (Biolegend), EPCR (eBio1560)(Invitrogen), IFITM1 (11727-3-AP)(ProteinTech), Caspase-1 (D-3)(Santacruz), Caspase-11 (8A5)(Novus), cleaved gasdermin (E3E3P)(Cell Signaling) and pSTAT1 (Tyr701, A17012A)(Biolegend). HSPC populations were identified with the following combinations: BM HSCs (CD48-CD150+CD34-Flt3-LSK), EPCR+ HSCs (CD48-CD150+EPCR+ LSK), CMP (CD16/32-CD34+ Lin-cKit+Sca1-(LKS-)), GMP (CD16/32+ LKS-), MEP (CD16/32-CD34-LKS-), CLP (CD16/32+Flt3+ LKS-), MkP (CD41+CD150+ LKS-) and CFUE (CD41-CD16/32-CD150-CD105+Ter119-LKS-) cells. IntraPrep (Beckman Coulter) was used prior to intracellular staining for pSTAT1 and cleaved gasdermin. Annexin V binding buffer (Biolegend) was used for staining Annexin V. FAM-FLICA (BIORAD) staining was conducted according to manufacturer protocol. BODIPY-SM (Invitrogen) was prepared according to manufacturer protocol but stained with modifications. Flow cytometric analysis and sorting was conducted on CytoFLEX (Beckman Coulter) and BD FACS Aria III cell sorter (BD Biosciences). Flow cytometric data was analyzed using FlowJo 10.1 (Tree Star).

### Complete blood counts

Peripheral blood was obtained from the superficial temporal vein of mice, collected in tubes containing EDTA, and analyzed using the Celltac Alpha veterinary hematology analyzer (Nihon Kohden).

### BM transplantation

BM MNCs (2×10^5^ cells) from C57BL/6-Ly5.1 mice together with 500 LT-HSCs from donor mice (Ly5.2) were transplanted into lethally-irradiated C57BL/6-Ly5.1 congenic mice. Secondary transplantations into lethally-irradiated C57BL/6-Ly5.1 congenic mice were performed using 2×10^6^ BM MNCs from primary recipients. PB donor chimerism was analyzed monthly. Data was recorded for 10000 to 30000 MNCs chimerism (depending on the recovery of the PB) for WBC. Recipient mice were sacrificed for analysis 4 months after BMT.

### Bulk RNA sequence, single-cell RNA sequence and ATAC sequence analysis

Bulk RNA sequence (GSE16700) and single-cell RNA sequence data (GSE320303) were obtained from previously reported data sets and re-analyzed ^17,25^. ATAC sequence analysis was conducted on HSCs obtained from *Thpo^+/+^*and *Thpo^-/-^* mice. Chromatin was extracted and processed for Tn5 transposase-mediated tagmentation and adapter incorporation for 30 min at 37 °C using the Nextera DNA Library Prep Kit (Illumina) according to the manufacturer’s protocol. Transposed DNA was amplified using NEBNext High-Fidelity 2× Master Mix (New England Biolabs). Samples were pooled and run on an Illumina HiSeq 4000 (Novogene). Reads that passed the quality filter step were mapped to the mouse reference genome (mm10) using Bowtie2 ^53^. Peaks were called jointly for replicates of each condition using Genrich. A consensus peak set was generated by merging peaks from both conditions, and reads within consensus peaks were quantified using featureCounts ^54^. Differentially accessible regions were identified from the read count matrix using DESeq2 ^55^. De novo motif discovery was performed using HOMER ^56^. Peaks were annotated to nearby genes, and the resulting gene lists were subjected to pathway enrichment analysis using ChIPseeker ^57^.

### Metabolomic (lipidome) analysis

Lipidome analysis was conducted according to Lipidome lab Non-targeted Lipidome Scan package (Lipidome lab, Akita, Japan), using liquid chromatograph orbitrap mass spectrometry (LC-OrbitrapMS) based on the methods described previously ^58,59^. Briefly, total lipids were extracted from cell samples with the modified Bligh-Dyer method. An aliquot of the lower/organic phase was evaporated to dryness under N_2_, and the residue was dissolved in methanol for LC-MS/MS measurements. Metabolite extraction and metabolomic analysis were conducted at Human Metabolome Technologies (HMT) (HMT, Tsuruoka, Yamagata, Japan).

### FRAP analysis

HSCs cultured in SFEM (Stem Cell Technologies) with Poly(I:C) were plated on a chambered coverglass (Matsunami) coated with Cell-Tak (Corning). HSCs were transiently stained in Dil stain (Thermo Fisher Scientific) prior to plating. FRAP analyses were performed following the protocols described previously ^60^ using the LSM-710 or LSM-900 with Airyscan2 confocal microscope with a water-immersion 60× lens of NA 1.2 at 37°C, and data were analyzed with Zen software. A small area (approximately a 1.0 μm diameter circle) was selected within the cell and bleached. Snapshots were then collected using as low laser power as 0.1% every 3-5 s.

### SEM analysis

HSCs were plated on coverslips coated with Cell-Tak (Corning) and rinsed with 0.1 M PBS and fixed in 2.5% glutaraldehyde and 2% paraformaldehyde in PBS for 1hr at 4℃. Cells were rinsed with PBS and fixed 1% osmium tetroxide (OsO_4_) in PBS for 1hr at 4℃. The samples were dehydrated through a graded ethanol series (50%, 70%, 90%, and 95% for 10 min each), followed by three 10-min changes of 100% ethanol and t-butyl alcohol freeze-dried using ID-2 (EIKO ENGINEERING) and were mounted onto an SEM stub and osmium-coated using NEOC (Meiwafosis). Images were obtained using a JSM-6610LA scanning electron microscope (JEOL).

### Statistical analysis

Statistical details of experiments can be found in Figure legends. All results are expressed as the mean±SD unless otherwise specified. Statistical significance was determined by two-tailed Student t-test, Tukey’s multiple comparison test and tests specified in figure legends. Outliers were assessed and removed using Grubb’s test. All experiments were repeated in at least two independent experiments.

---

**Extended data 1:**
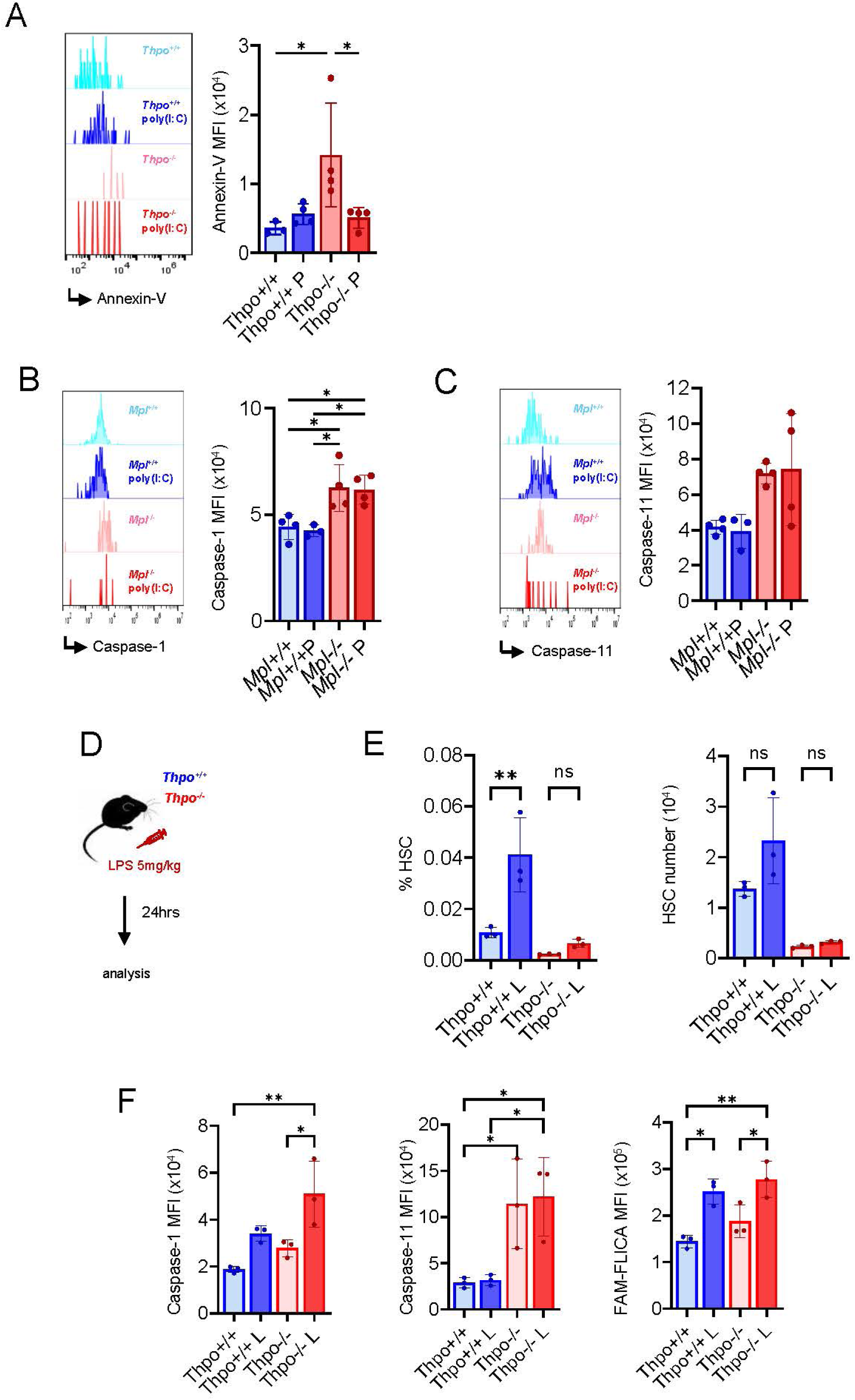
(A) Representative flow cytometric plots and quantification of Annexin-V staining on HSCs from *Thpo^+/+^* and *Thpo^-/-^* mice with or without poly(I:C) (P). (mean±SD) (n>3) * P<0.05 by Tukey test. Representative flow cytometric plots and quantification of Caspase-2 (B), caspase-11 (C) staining on HSCs from *Mpl^+/+^* and *Mpl^-/-^* mice with or without poly(I:C) (P). (mean±SD) (n>3) * P<0.05, ** P<0.01 and *** P<0.001 by Tukey test. (D) Schematic representation of LPS administration to *Thpo^+/+^* and *Thpo^-/-^* mice. (E) HSC frequency and number from *Thpo+/+* and *Thpo^-/-^* mice with or without LPS (L) administration. (mean±SD) (n=3) ** P<0.01, ns: not significant by Tukey test. (F) Representative flow cytometric plots and quantification of Caspase-2, caspase-11 and FAM-FLICA staining on HSCs from *Thpo+/+* and *Thpo^-/-^* mice with or without LPS (L) administration. (mean±SD) (n=3) * P<0.05 and ** P<0.01 by Tukey test.

**Extended data 2:**
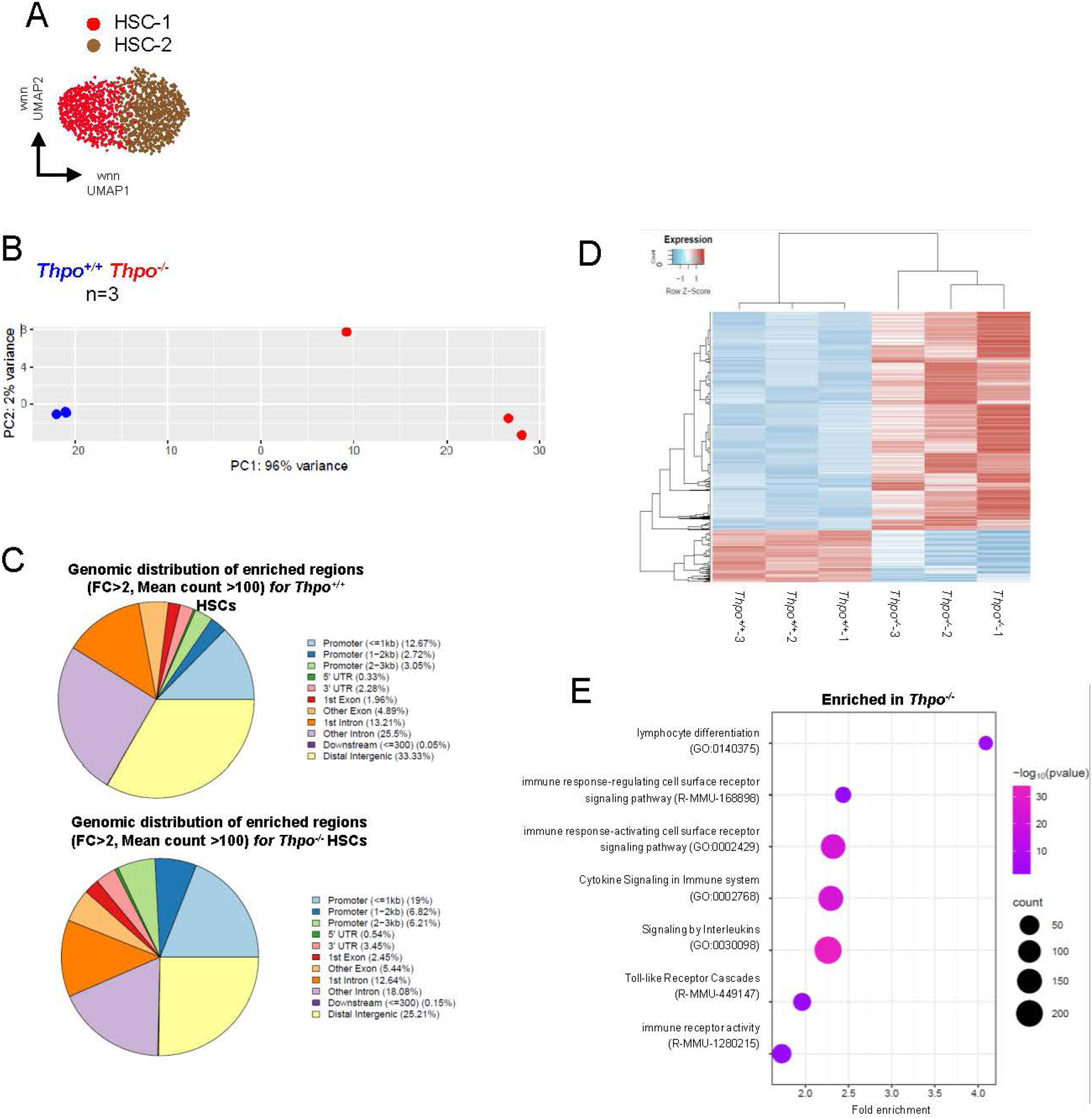
(A) UMAP showing HSC-1 and HSC-2 clusters. (B) PCA plots for ATAC-seq data conduced on *Thpo^+/+^*and *Thpo^-/-^*HSCs. (C) Genomic distribution of enriched regions for *Thpo^+/+^* and *Thpo^-/-^*HSCs. (D) Heatmap showing significantly accessible chromatin regions in *Thpo^+/+^*and *Thpo^-/-^*HSCs. (E) Gene ontology analysis of genes enriched in *Thpo^-/-^*HSCs.

**Extended data 3:**
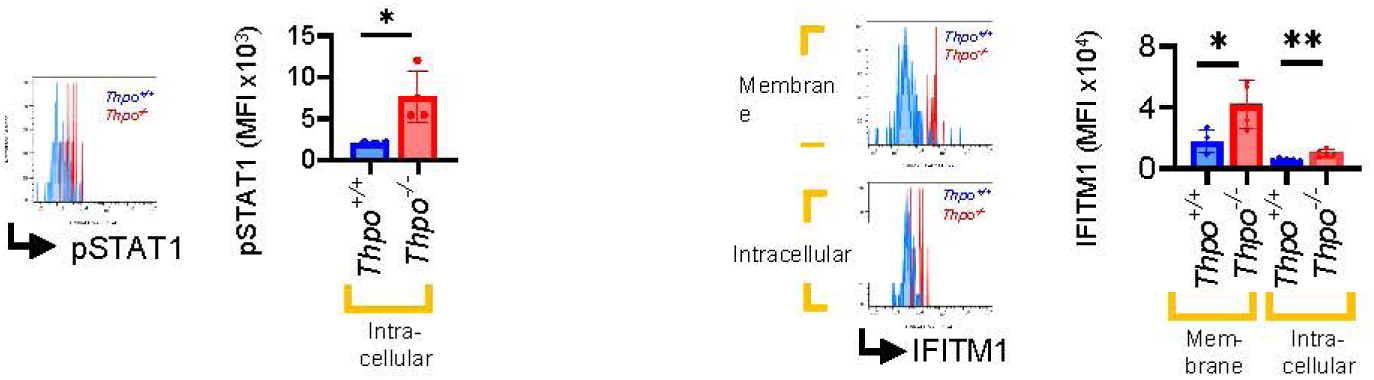
Representative flow cytometric plots and quantification of intracellular expression of pSTAT1 and membrane and intracellular expression of IFITM1 on *Thpo^+/+^* and *Thpo^-/-^*HSCs. (mean±SD) (n=4) * P<0.05, ** P<0.01 by Student t-test.

**Extended data 4:**
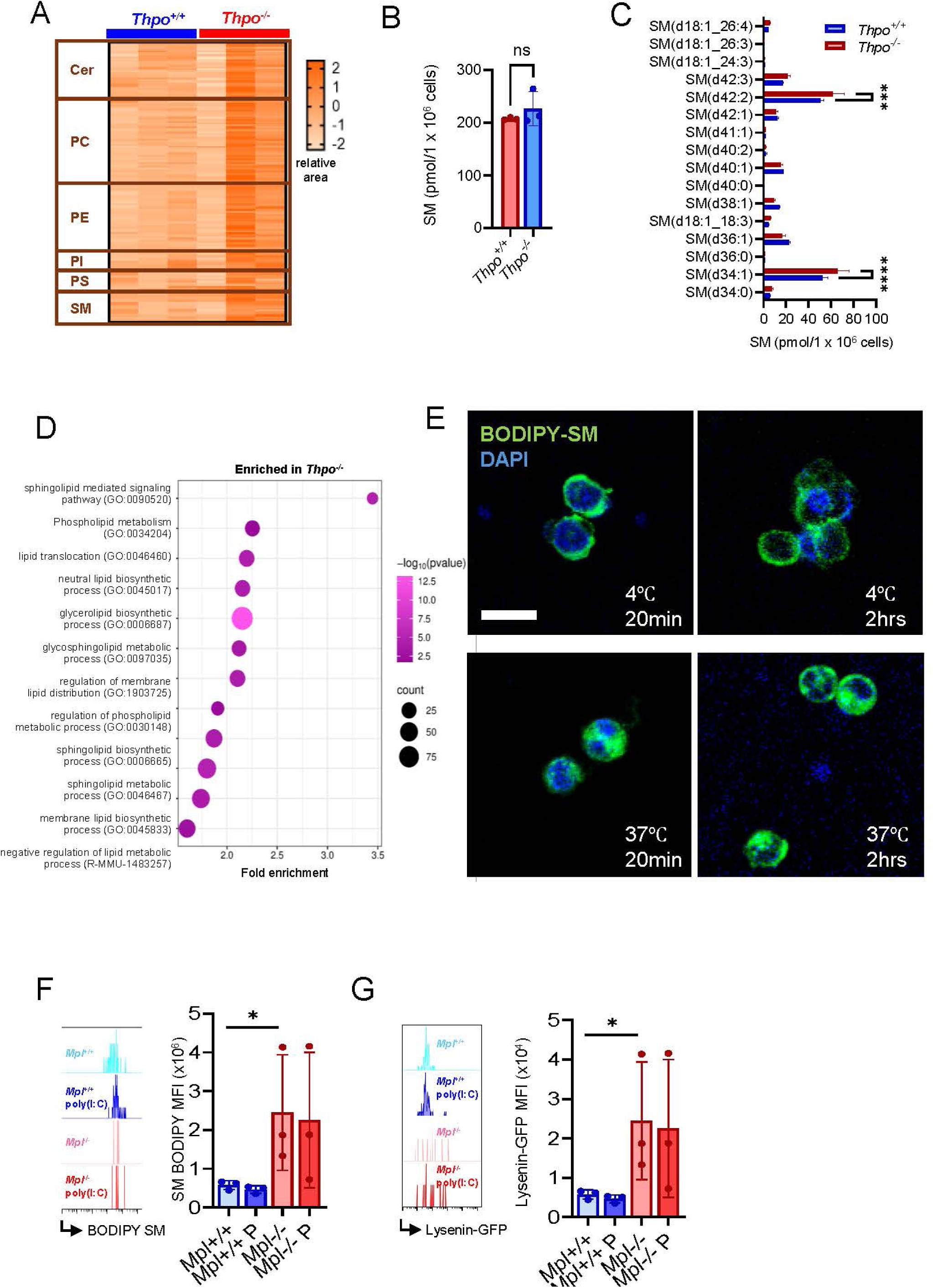
(A) Heatmap of relative area values of various cell membrane lipids (sphingomyelin (SM), phosphatidylserine (PS), phosphatidylinositol (PI), phosphatidylethanolamine (PE), phosphatidylcholine (PC) and ceramide (Cer)) expressed in *Thpo^+/+^*and *Thpo^-/-^* HSCs. (B) Total SM concentration of HSCs from *Thpo^+/+^* and *Thpo^-/-^* mice. ns: not significant by Student t-test. (C) Quantification of SMs in *Thpo^-/-^* and *Thpo^+/+^* HSCs. (mean±SD) (n=3) *** P<0.001, **** P<0.0001 by Student t-test. (D) Gene ontology analysis of genes enriched in *Thpo^-/-^*HSCs from ATAC-seq. (E) Fluorescent microscopic images of HSCs stained with BODIPY-SM under indicated conditions. Note that staining at 4℃ shows probe accumulation on cell membrane while the probe is incorporated into the cell when stained at 37℃. Scale bar = 10μm. (F) Representative flow cytometric plot and MFI of BODIPY-SM staining on cell membrane in HSCs from *Mpl^+/+^*and *Mpl^-/-^* mice treated with poly(I:C). (mean±SD) (n>3) * P<0.05 by Tukey test. (G) Representative flow cytometric plot and MFI of Lysenin-GFP staining on cell membrane in HSCs from *Mpl^+/+^* and *Mpl^-/-^* mice treated with poly(I:C). (mean±SD) (n>3) * P<0.05 by Tukey test.

**Extended data 5:**
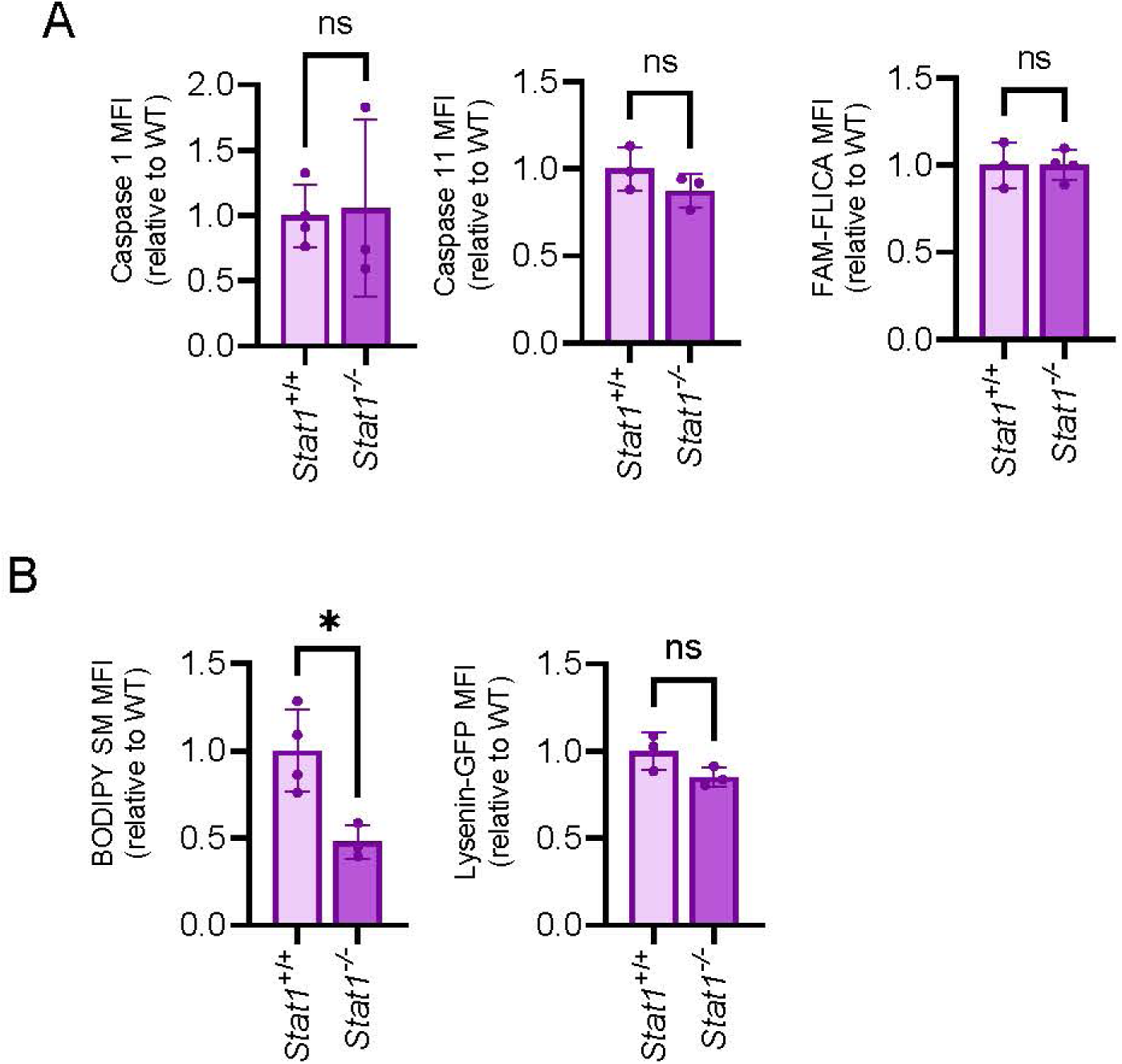
(A) MFI of expression of caspase-1, caspase-11 and active caspase-1 (FAM- FLICA) staining on HSCs from *Stat1^+/+^*and *Stat1^-/-^* mice. Values are shown relative to WT. (mean±SD) (n>3) ns: not significant by Student t-test. (B) MFI of BODIPY- SM and Lysenin-GFP staining on cell membrane in HSCs from *Stat1^+/+^* and *Stat1^-/-^* mice. Values are shown relative to WT. (mean±SD) (n>3) * P<0.05 and ns: not significant by Student t-test.

## Notes

### Competing Interest Statement

The authors have declared no competing interest.

## References

1. Rettkowski, J. & Cabezas-Wallscheid, N. Regulation of Hematopoietic Stem Cell Dormancy and Quiescence: Insights into Regeneration and Disease. Annu. Rev. Cell Dev. Biol. 41, 231–258 (2025).

2. Nakamura-Ishizu, A., Takizawa, H. & Suda, T. The analysis, roles and regulation of quiescence in hematopoietic stem cells. Development 141, 4656–4666 (2014).

3. Arai, F. et al. Tie2/angiopoietin-1 signaling regulates hematopoietic stem cell quiescence in the bone marrow niche. Cell 118, 149–61 (2004).

4. Passegué, E., Wagers, A. J., Giuriato, S., Anderson, W. C. & Weissman, I. L. Global analysis of proliferation and cell cycle gene expression in the regulation of hematopoietic stem and progenitor cell fates. J. Exp. Med. 202, 1599–611 (2005).

5. Essers, M. A. G. et al. IFNalpha activates dormant haematopoietic stem cells in vivo. Nature 458, 904–8 (2009).

6. Bogeska, R. et al. Inflammatory exposure drives long-lived impairment of hematopoietic stem cell self-renewal activity and accelerated aging. Cell Stem Cell 29, 1273–1284.e8 (2022).

7. Caiado, F., Pietras, E. M. & Manz, M. G. Inflammation as a regulator of hematopoietic stem cell function in disease, aging, and clonal selection. J. Exp. Med. 218, (2021).

8. Zeng, A. G. X. et al. Human haematopoietic stem cells remember inflammatory stress. Nature 2026 1–10 (2026) doi:10.1038/s41586-026-10522-7.

9. Wilson, A. et al. Dormant and self-renewing hematopoietic stem cells and their niches. Ann. N. Y. Acad. Sci. 1106, 64–75 (2007).

10. Kaushansky, K. Thrombopoietin. N. Engl. J. Med. 339, 746–54 (1998).

11. Gurney, A. L., Carver-Moore, K., de Sauvage, F. J. & Moore, M. W. Thrombocytopenia in c-mpl-deficient mice. Science 265, 1445–7 (1994).

12. de Sauvage, F. J. et al. Stimulation of megakaryocytopoiesis and thrombopoiesis by the c-Mpl ligand. Nature 369, 533–8 (1994).

13. Yoshihara, H. et al. Thrombopoietin/MPL signaling regulates hematopoietic stem cell quiescence and interaction with the osteoblastic niche. Cell Stem Cell 1, 685–97 (2007).

14. Qian, H. et al. Critical role of thrombopoietin in maintaining adult quiescent hematopoietic stem cells. Cell Stem Cell 1, 671–84 (2007).

15. de Laval, B. et al. Thrombopoietin-increased DNA-PK-dependent DNA repair limits hematopoietic stem and progenitor cell mutagenesis in response to DNA damage. Cell Stem Cell 12, 37–48 (2013).

16. de Laval, B. et al. Thrombopoietin promotes NHEJ DNA repair in hematopoietic stem cells through specific activation of Erk and NF-κB pathways and their target IEX-1. Blood blood-2013-07-515874-(2013) doi:10.1182/blood-2013-07-515874.

17. Nakamura-Ishizu, A. et al. Prolonged maintenance of hematopoietic stem cells that escape from Thrombopoietin deprivation. Blood https://doi.org/10.1182/blood.2020005517 (2021) doi:10.1182/blood.2020005517.

18. Mochizuki-Kashio, M., et al. Hematopoietic stem and progenitor cell hierarchy is established by thrombopoietin-driven neonatal hematopoiesis. Stem Cell Reports 21, (2026).

19. Ivashkiv, L. B. & Donlin, L. T. Regulation of type I interferon responses. Nat. Rev. Immunol. 14, 36 (2014).

20. Hu, X. & Ivashkiv, L. B. Cross-regulation of signaling pathways by interferon-gamma: implications for immune responses and autoimmune diseases. Immunity 31, 539–550 (2009).

21. Mitroulis, I. et al. Modulation of Myelopoiesis Progenitors Is an Integral Component of Trained Immunity. Cell 172, 147–161.e12 (2018).

22. Kaufmann, E. et al. BCG Educates Hematopoietic Stem Cells to Generate Protective Innate Immunity against Tuberculosis. Cell 172, 176–190.e19 (2018).

23. Walter, D. et al. Exit from dormancy provokes DNA-damage-induced attrition in haematopoietic stem cells. Nature 520, 549–52 (2015).

24. Haas, S. et al. Inflammation-Induced Emergency Megakaryopoiesis Driven by Hematopoietic Stem Cell-like Megakaryocyte Progenitors. Cell Stem Cell 17, 422–34 (2015).

25. Mochizuki-Kashio, M., et al. Hematopoietic stem and progenitor cell hierarchy is established by thrombopoietin-driven neonatal hematopoiesis. Stem Cell Reports 21, (2026).

26. Narayana, S. K. et al. The Interferon-induced Transmembrane Proteins, IFITM1, IFITM2, and IFITM3 Inhibit Hepatitis C Virus Entry. J. Biol. Chem. 290, 25946–25959 (2015).

27. Li, K. et al. IFITM proteins restrict viral membrane hemifusion. PLoS Pathog. 9, (2013).

28. Wilt, I. et al. IFITM1 and IFITM3 cooperate to restrict virus entry in endolysosomes. bioRxiv https://doi.org/10.1101/2025.06.01.657267 (2025) doi:10.1101/2025.06.01.657267.

29. Westerterp, M. et al. Cholesterol Accumulation in Dendritic Cells Links the Inflammasome to Acquired Immunity. Cell Metab. 25, 1294–1304.e6 (2017).

30. Wei, X. et al. Fatty acid synthesis configures the plasma membrane for inflammation in diabetes. Nature 539, 294–298 (2016).

31. Zhou, Q. D. et al. Interferon-mediated reprogramming of membrane cholesterol to evade bacterial toxins. Nat. Immunol. 21, 746–755 (2020).

32. Molenaar, M. R. et al. LION/web: a web-based ontology enrichment tool for lipidomic data analysis. Gigascience 8, 1–10 (2019).

33. Ramstedt, B. & Slotte, J. P. Membrane properties of sphingomyelins. FEBS Lett. 531, 33–37 (2002).

34. Endapally, S. et al. Molecular Discrimination between Two Conformations of Sphingomyelin in Plasma Membranes. Cell 176, 1040–1053.e17 (2019).

35. Ramstedt, B. & Slotte, J. P. Membrane properties of sphingomyelins. FEBS Lett. 531, 33–37 (2002).

36. Ishitsuka, R. & Kobayashi, T. Lysenin: a new tool for investigating membrane lipid organization. Anat. Sci. Int. 79, 184–190 (2004).

37. Barbieri, D. et al. Thrombopoietin protects hematopoietic stem cells from retrotransposon-mediated damage by promoting an antiviral response. J. Exp. Med. 215, 1463 (2018).

38. Bruns, I. et al. Megakaryocytes regulate hematopoietic stem cell quiescence through CXCL4 secretion. Nat. Med. https://doi.org/10.1038/nm.3707 (2014) doi:10.1038/nm.3707.

39. Zhao, M. et al. Megakaryocytes maintain homeostatic quiescence and promote post-injury regeneration of hematopoietic stem cells. Nat. Med. https://doi.org/10.1038/nm.3706 (2014) doi:10.1038/nm.3706.

40. Nakamura-Ishizu, A., Takubo, K., Fujioka, M. & Suda, T. Megakaryocytes are essential for HSC quiescence through the production of thrombopoietin. Biochem. Biophys. Res. Commun. 454, 353–357 (2014).

41. Matatall, K. A., Shen, C. C., Challen, G. A. & King, K. Y. Type II interferon promotes differentiation of myeloid-biased hematopoietic stem cells. Stem Cells 32, 3023–3030 (2014).

42. Li, J. et al. STAT1 is essential for HSC function and maintains MHCIIhi stem cells that resist myeloablation and neoplastic expansion. Blood 140, 1592– 1606 (2022).

43. Sanjuan-Pla, A. et al. Platelet-biased stem cells reside at the apex of the haematopoietic stem-cell hierarchy. Nature 502, 232–236 (2013).

44. Nakamura-Ishizu, A. et al. Thrombopoietin Metabolically Primes Hematopoietic Stem Cells to Megakaryocyte-Lineage Differentiation. Cell Rep. 25, 1772–1785.e6 (2018).

45. Alvarado, L. J. et al. Eltrombopag maintains human hematopoietic stem and progenitor cells under inflammatory conditions mediated by IFN-γ. Blood 133, 2043–2055 (2019).

46. Kao, Y.-R. et al. Thrombopoietin receptor–independent stimulation of hematopoietic stem cells by eltrombopag. Sci. Transl. Med. 10, eaas9563 (2018).

47. Swanson, K. V., Deng, M. & Ting, J. P. Y. The NLRP3 inflammasome: molecular activation and regulation to therapeutics. Nat. Rev. Immunol. 19, 477–489 (2019).

48. Yilmaz, N., Yamaji-Hasegawa, A., Hullin-Matsuda, F. & Kobayashi, T. Molecular mechanisms of action of sphingomyelin-specific pore-forming toxin, lysenin. Semin. Cell Dev. Biol. 73, 188–198 (2018).

49. Pedrera, L. et al. The Important Role of Membrane Fluidity on the Lytic Mechanism of the α-Pore-Forming Toxin Sticholysin I. Toxins (Basel). 15, (2023).

50. Kimura, S., Roberts, A. W., Metcalf, D. & Alexander, W. S. Hematopoietic stem cell deficiencies in mice lacking c-Mpl, the receptor for thrombopoietin. Proc. Natl. Acad. Sci. U. S. A. 95, 1195–200 (1998).

51. Meraz, M. A. et al. Targeted disruption of the Stat1 gene in mice reveals unexpected physiologic specificity in the JAK-STAT signaling pathway. Cell 84, 431–442 (1996).

52. Takizawa, H., Regoes, R. R., Boddupalli, C. S., Bonhoeffer, S. & Manz, M. G. Dynamic variation in cycling of hematopoietic stem cells in steady state and inflammation. J. Exp. Med. 208, 273–84 (2011).

53. Langmead, B. & Salzberg, S. L. Fast gapped-read alignment with Bowtie 2. Nat. Methods 9, 357–9 (2012).

54. Liao, Y., Smyth, G. K. & Shi, W. featureCounts: an efficient general purpose program for assigning sequence reads to genomic features. Bioinformatics 30, 923–930 (2014).

55. Love, M. I., Huber, W. & Anders, S. Moderated estimation of fold change and dispersion for RNA-seq data with DESeq2. Genome Biol. 15, (2014).

56. Heinz, S. et al. Simple combinations of lineage-determining transcription factors prime cis-regulatory elements required for macrophage and B cell identities. Mol. Cell 38, 576–589 (2010).

57. Yu, G., Wang, L. G. & He, Q. Y. ChIPseeker: an R/Bioconductor package for ChIP peak annotation, comparison and visualization. Bioinformatics 31, 2382–2383 (2015).

58. Takumi, H. et al. Comprehensive Analysis of Lipid Composition in Human Foremilk and Hindmilk. J. Oleo Sci. 71, 947–957 (2022).

59. Nishiumi, S. et al. Comparative Evaluation of Plasma Metabolomic Data from Multiple Laboratories. Metabolites 12, (2022).

60. Chen, C. et al. Phase separation and toxicity of c9orf72 poly(Pr) depends on alternate distribution of arginine. Journal of Cell Biology 220, (2021).

